# On-cell Saturation Transfer Difference (STD) NMR on ion channels: characterizing negative allosteric modulator binding interactions of P2X7

**DOI:** 10.1101/2025.04.22.649959

**Authors:** Serena Monaco, Jacob Browne, Matthew Wallace, Jesus Angulo, Leanne Stokes

## Abstract

P2X7 receptors are important drug targets involved in pathologies ranging from psychiatric disorders to cancer. Being membrane embedded receptors, they are challenging for structural characterisation and at present we only have a small number of X-ray and cryoEM structures for P2X7 bound to antagonists. We demonstrate that Saturation Transfer Difference (STD) NMR on live mammalian cells (on-cell STD NMR) overexpressing P2X7 receptors allows further structural insight on the complexes of P2X7 with two potent negative allosteric modulators, namely AZ10606120 and JNJ-47465567, via the determination of the binding epitope mapping of the interactions e.g. the main region of contact between the ligand and the binding pocket. This approach, reported for the first time on membrane-embedded ion channels, in combination with molecular docking, allows us to propose the first NMR-validated 3D molecular models for two antagonists as bound to human P2X7 receptors, and to correlate the structural knowledge acquired with the pharmacology data. We highlight the transformative potential of this application to aid drug design efforts in a less resource-demanding fashion than X-ray crystallography and cryo-EM and we envisage on-cell STD NMR to fast become an asset for structure-activity-relationship studies helping knowledge-based development of efficient drugs targeting P2X7 and other ion channels/membrane-embedded proteins.

## Introduction

P2X7 receptors are important ligand-gated ion channels highlighted as therapeutic targets for psychiatric disorders such as depression [1], autism-spectrum disorders [2, 3], and cancer [4]. Ligand-gated ion channels respond to chemical molecules that interact with specific binding pockets on the channel complex. Once bound to the orthosteric site, agonist molecules induce conformational changes to the protein complex allowing intrinsic ion channel pore opening. Allosteric sites also exist on ion channels which can either hinder or enhance the ion channel opening mechanisms. X-ray crystallography has identified three identical inter-subunit negative allosteric modulator (NAM) sites on P2X7 capable of accepting a diverse array of chemicals, as demonstrated by crystal structures of five antagonists in complex with giant panda P2X7 [5]. These sites are located behind the three orthosteric ATP sites, where the allosteric modulator can hinder ATP-driven conformational changes by acting as molecular wedges [5].

Many antagonists have been developed for P2X7 by multiple pharmaceutical companies and academic groups [6]. Several P2X7 antagonists have progressed into clinical trials including AZD9056 [7, 8] and CE-224,535 [9] for rheumatoid arthritis and Crohn’s disease, GSK1482160 (Phase I) [10], and JNJ-54175446 (current Phase II trials for depressive disorders) [11] although none have yet been taken further. Some pharmacological studies have used NAM binding site mutagenesis of hP2X7 to confirm the involvement of the NAM site in antagonist effects [12, 13] however, as yet, there is no published crystal or cryo-EM structure of human P2X7 in complex with antagonists. Furthermore, some antagonists display large species-dependent differences in activity whilst others display only subtle differences in potency.

In the effort to advance the structural knowledge on human P2X7 and overcome the severe limitations of X-ray and cryo-EM to study membrane embedded receptors, we have used saturation transfer difference (STD) NMR spectroscopy to study NAM antagonists binding to interspecies P2X7 receptors in solution [14]. To the best of our knowledge, this is first example of the use of STD NMR to study membrane-embedded ion channels on living cells. Traditionally, STD NMR [15] has been used to study protein-ligand interactions in solution. It relies on the selective saturation of the protein, which is transferred to the ligand, causing a reduction of the ligand NMR signal intensities. The STD NMR spectra are obtained as the difference of the reference NMR spectra (where selective saturation is off-resonance) minus the on-resonance irradiated NMR spectra. Importantly, the STD intensity for each ligand proton is quantitatively related to the proximity of the given ligand proton to the protein surface. This allows a determination of the so-called binding epitope, i.e. a map depicting the ligand protons in closer contact to the protein, informing on the ligand binding mode in the protein binding pocket (Figure 1) [16]. This ligand-based NMR technique has been previously proposed to investigate ligand binding to an over-expressed receptor on the surface of live cells (rather than purified in solution) [17]. In this way, the determination of ligand-protein interactions/contacts with the protein attached to the cell membrane or embedded within it are achieved, without the need for sample manipulations, labels, or protein purification. This confers strong potential to the approach, as it can provide structural information about the biomolecular interactions in conditions close to their native-like environment.

**Figure 1:**
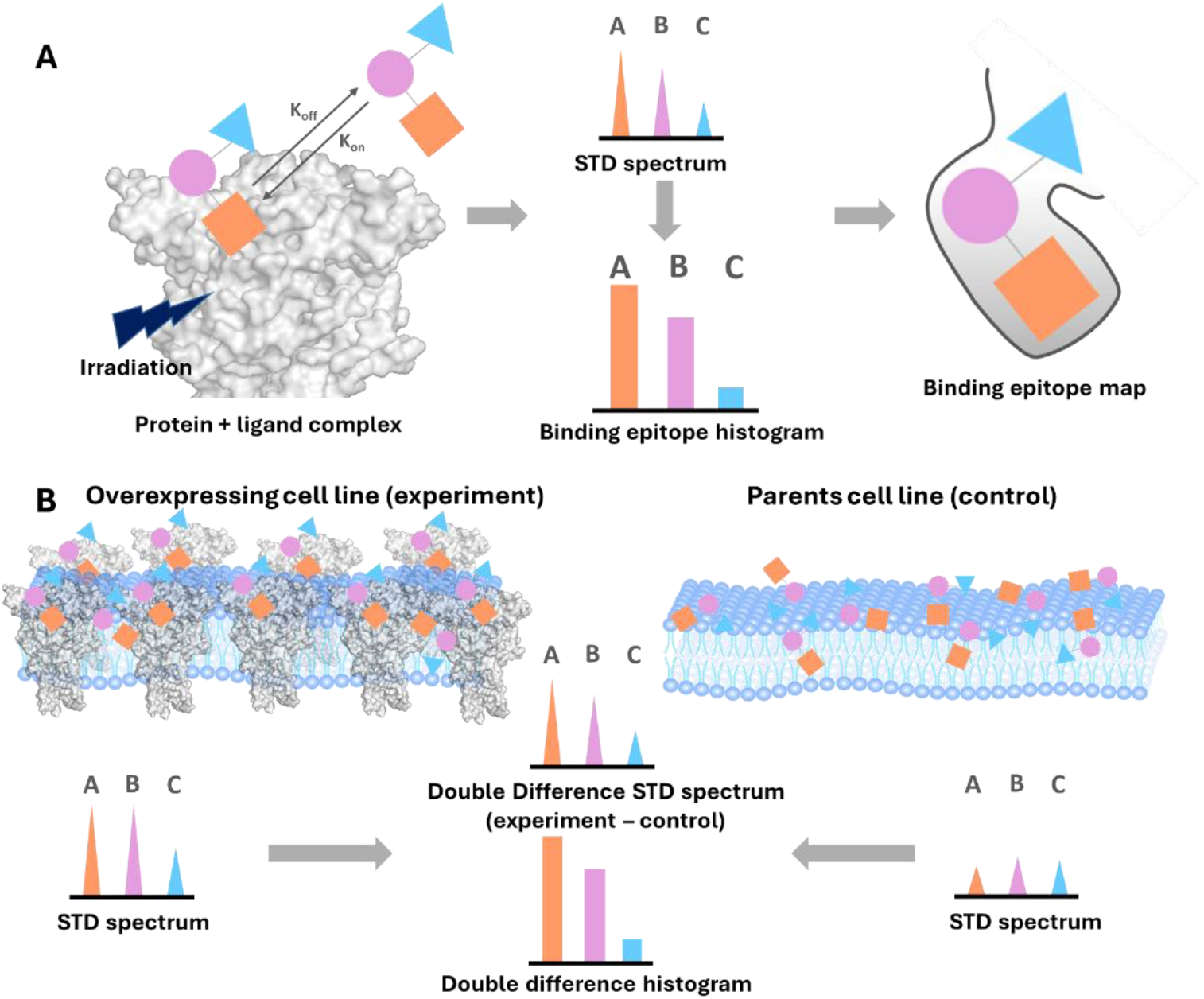
Principles behind on-cell STD NMR. **A:** representation of binding epitope map determination by STD NMR. **B:** representation of double-difference STD NMR, a necessary variation of the technique for on-cell STD NMR experiments, to eliminate signals resulting from the ligand interacting with the cell membrane.

To ensure cancellation of signals from residual ligand interactions with cell membranes or other proteins on the cell surface, it is necessary to carry out a double-difference (STDD NMR) version of the experiment, carrying out the subtraction between the STD spectra obtained in the presence of the “positive” cell line overexpressing the receptor of interest, minus the STD spectra obtained with the “parent” cell line or control experiment [17]. This isolates, for each set of protons, the actual STD response coming from the specific interaction of the small molecule with the overexpressed receptor. Following pioneering work from Jimenez-Barbero and co-workers on DC-SIGN interactions [18], several applications have exploited this approach, extending the technique from sole binding screening to on-cell STD binding epitope mapping [19-21]. A large part of these studies has focused on protein-glycan interactions, given the many crucial processes involving cell-pathogen, cell-cell, and cell-matrix interactions, mostly taking place through surface proteins.

In this work, we have applied on-cell STD NMR to determine small molecule ligand binding epitope mapping for *membrane-embedded* P2X7 ion channel overexpressed in HEK-293 cells. We have characterised the interactions of P2X7 with two well studied NAMs, AZ10606120 and JNJ-47965567, for which crystal structures are available for the related giant panda P2X7 (pdb entries: 5u1w, 5u1x). To understand the effect on the ligand-protein interactions upon mutation of the NAM site, we analysed P2X7 from different species (human, mouse and rat) as well as human P2X7 carrying mutations in the NAM pocket. We generated clonal cell lines expressing three signature P2X7 NAM-site mutations (F88A, M105A and F103A) [5, 12] and used computational docking to propose 3D molecular models of the antagonists AZ10606120 and JNJ-47965567 binding at human, rat and mouse P2X7 homology models. We then validated these against the on-cell STD NMR data using a recently disclosed reduced relaxation matrix approach (RedMat) that allows time-efficient prediction of STD binding epitope mappings from 3D molecular models [22].

Our approach provides, for the first time, structural insights on the human and rodent P2X7 receptors in complex with NAM antagonists. We also demonstrate that binding orientations determined by STD NMR can be correlated with pharmacological data and thus understand how differences in potency arise between different allosteric modulators across species and mutants.

## Methods

### Cell culture

Genetically edited HEK-293 cells lacking P2Y2 receptors were used to generate stable cell lines expressing human P2X7 (hP2X7), mouse P2X7 (mP2X7), rat P2X7 (rP2X7) or mutants of hP2X7 NAM site (F88A, M105A and F103A). Transfection was performed using lipofectamine 2000 (Fisher Scientific) and cells were subjected to geneticin (800 µg/ml) selection for 2-3 weeks. This was followed by single cell cloning by limiting dilution in 96-well plates. Individual clones were selected for further study based on ATP-induced TO-PRO-3 uptake on a BD Cytoflex flow cytometer. Cell lines were maintained in DMEM:F12 media supplemented with 10% foetal bovine serum (PanBiotech), penicillin and streptomycin (Fisher Scientific) and kept in a humidified incubator with 5% CO_2_. Stable P2X7-expressing cells were kept under geneticin selection at 400 μg/ml. Passaging was performed twice weekly using 0.25% Trypsin-EDTA (Fisher Scientific).

### Intracellular calcium measurements

P2X7 responses were measured by monitoring intracellular calcium responses using fura-2AM loaded cells. Cells were plated at 2-2.5 x 10^4^ cells/well the day before experiments in 96-well plates (NUNC) coated with 50 μg/ml poly-D-lysine (Merck Millipore). A loading buffer containing 2 μM fura-2AM (HelloBio) and 250 μM sulfinpyrazone (Merck Millipore) in a low divalent buffer (145mM NaCl, 5mM KCl, 0.2mM CaCl_2_, 10mM HEPES, 13mM glucose, pH 7.3) was added to cells for 45 minutes at 37°C. After loading, the buffer was removed and replaced with calcium-containing extracellular solution (145mM NaCl, 5mM KCl, 2mM CaCl_2_, 10mM HEPES, 13mM glucose, pH 7.3). Responses were measured using a Flexstation 3 plate reader with excitation wavelengths 340nm and 380nm and single emission wavelength 510nm. Sample interval was 3.5 seconds and each well was recorded for 180 seconds. Fura-2 ratios were calculated in Softmax Pro v5.4, zero baseline was applied and responses were measured as normalised area under curve (AUC). Dose inhibition curves were plotted using GraphPad Prism v6 non-linear regression (three parameter fit) and logIC_50_ values reported.

### Cell preparation for STD NMR

Cells were grown in 10 cm tissue culture petri dishes (Greiner) to confluency in DMEM:F12 complete media containing 400 μg/ml geneticin. 0.25% trypsin-EDTA was used to dissociate cells and fresh media used to inactivate trypsin. Cells were then washed twice in PBS using centrifugation at 350g and washed into deuterated-PBS (1ml). Cells were resuspended in 450 μl deuterated-PBS and carefully counted using a hemocytometer. For each NMR experiment 2 million cells in a total volume of 450 μl was used and ligand was added to 0.3 mM concentration. For JNJ-47965567, DMSO-d6 was added to aid solubility (40 μl) and the volume of deuterated-PBS adjusted accordingly.

### Antagonist NMR assignment and STD NMR analysis

All the NMR experiments were recorded at ^1^H frequency of 800.23 MHz with a Bruker Avance III spectrometer equipped with a 5-mmD probe TXI 800 MHz H-C/N-D-05 Z BTO. Firstly, the antagonists AZ10606120 and JNJ-47665567 were analysed by 1D and 2D NMR in their free state and assigned using standard ^1^H-^1^H COSY (cosydfesgpph) and ^1^H-^13^C HSQC (hsqctgpsp) experiments, at 293K.

For STD NMR experiments, the samples were prepared as described in the above section, the temperature was set at 293 K and the NMR tubes were kept spinning in the probe for the entire duration of the experiment to ensure that the cells stayed well in suspension. An STD pulse sequence that included 2.5 ms and 5 ms trim pulses and a 3 ms spoil gradient and water suppression by excitation sculpting with gradients was used (*stddiffesgp*.*3*). Saturation was achieved by applying a train of 50 ms Gaussian pulses (0.40 mW) on the f2 channel, at 0 ppm (on-resonance experiments) and 40 ppm (off-resonance experiments). The broad protein signals were removed by using a 40 ms spinlock (T_1ρ_) filter. The experiments were performed at the single saturation time of 2 s, with a d1 delay of 3 s, with 4 dummy scans and 192 scans, to keep the total duration of the experiment to 35 minutes. Control experiments with a 6 s d1 delay were performed to make sure that the d1 did not affect the binding epitope. STD NMR experiments were performed in duplicate for each ligand in the presence of each, positive and negative parent cell lines. Biological replicates were performed from distinct cell culture plates on different days.

Binding epitopes were obtained by determining the STD% for each ligand signal (corresponding to one or more protons, depending on signal overlapping). The analysis was performed for the “positive” cell lines and the parent “negative” cell line separately and the double difference was performed on the processed STD%, for each biological duplicate. The double differences were then averaged between the duplicates and normalised to the most intense one for each complex (for which 100% is arbitrarily assigned). Error bars on the normalised STD percentage bar charts have been calculated by determining the standard deviation of the raw STD values for each atom (before normalisation) and then normalising each standard deviation to the percentage STD value.

### Computational docking and analysis

Homology models for human P2X7 and mouse P2X7 were prepared using SWISS model and template structure 5u1l. The cryoEM structure of rat P2X7 (PDB 6u9v) was truncated at amino acid residue 401 and minimised in Maestro using OPLS4 forcefield. Similar relaxation was performed for hP2X7 and mP2X7 models. Template-based docking was performed using CBdock2 (https://cadd.labshare.cn/cb-dock2/index.php), a web-accessible version of Autodock Vina, for AZ10606120 and JNJ-47965567 aligning with database templates such as 5u1l. Only one docking pose was generated using this approach.

### RedMat

Docking poses were reviewed using RedMat (http://redmat.iiq.us-csic.es/.) which uses a reduced relaxation matrix approach to predict binding epitope mappings of ligands in complex with a receptor protein. Docking poses were first processed by removing atom charges and renumbering ligand atoms as to prevent any numerical overlap with the receptor protons. A complex correlation time of 50 ns was used together with a ligand concentration of 0.3 mM, a receptor concentration of 50 µM, a complex dissociation constant of 0.01 µM and a cut-off distance of 12 Å. Methyl protons were selected to be selectively irradiated and the spectrometer ^1^H frequency used was 800MHz. R-NOE factors were calculated following the equation for each set of experimental and predicted data.

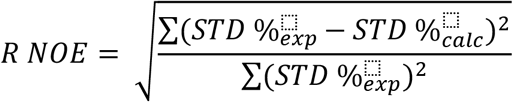

## Results

To perform the on-cell STD NMR experiments we used HEK-293 cells over-expressing hP2X7 as “positive” cell line, and non-expressing HEK-293 cells as the “negative” cell line (control background) for subtraction. We established a protocol involving several washes to remove traces of media and to equilibrate cells into deuterated-PBS. A careful cell count was critical and the number of cells in the positive and the negative cell line were kept equivalent with 2 million cells routinely used. Once the ligand was added at a concentration of 0.3 mM, the cells were placed into the spectrometer (800 MHz) and STD NMR experiments were performed at a single saturation time (2 s), as described in Methods. Critically, each experiment lasted 35 minutes and was performed while spinning the NMR sample tube to ensure the cells were well suspended in the buffer.

### On-cell STD NMR and docking studies on WT hP2X7 in complex with AZ10606120 and JNJ JNJ-47965567

In Figure 2A, the 1D NMR spectra for AZ10606120 are shown together with the assignment of protons. STD% were extracted for each set of protons in the positive and the negative samples, and then their values were subtracted. The proton with the highest STD was arbitrarily assigned a 100% value in the epitope mapping process and the remaining intensities were normalised relative to that. Figure 2B shows a histogram depicting the binding epitope of AZ10606120 (each bar corresponds to the normalised STD intensities of each given proton set with error bars displaying variability between replicates). Two ligand binding regions were clearly identified: a “core” region with strong saturation transfer (the quinoline moiety) and two areas with weaker saturation transfer, i.e. the aliphatic tail (“a” “b” “c” and “d” in Figure 2B) and the adamantane moiety (“Ad” in Figure 2B). We compared the ligand pose in the crystal structure of AZ10606120 at pdP2X7 (pdb:5u1w) and the result from the computational docking of AZ10606120 to a human P2X7 homology model (Figure 2C) and used RedMat [22] to correlate the experimental on-cell STD NMR binding epitope data to the predicted computational pose. RedMat simulates the ligand binding epitope mapping from a 3D molecular model of a protein-ligand complex and compares it with the experimental data. Then, the so-called R-NOE factor evaluate the agreement between the predicted and the experimentally determined epitope map, where an R-NOE close to 0.3 is typically considered as indicative of good agreement, and hence supporting experimental validation of the proposed 3D model. The R-NOE value for AZ10606120 as bound to hP2X7 was 0.22, suggesting excellent correlation with the experimental data.

**Figure 2:**
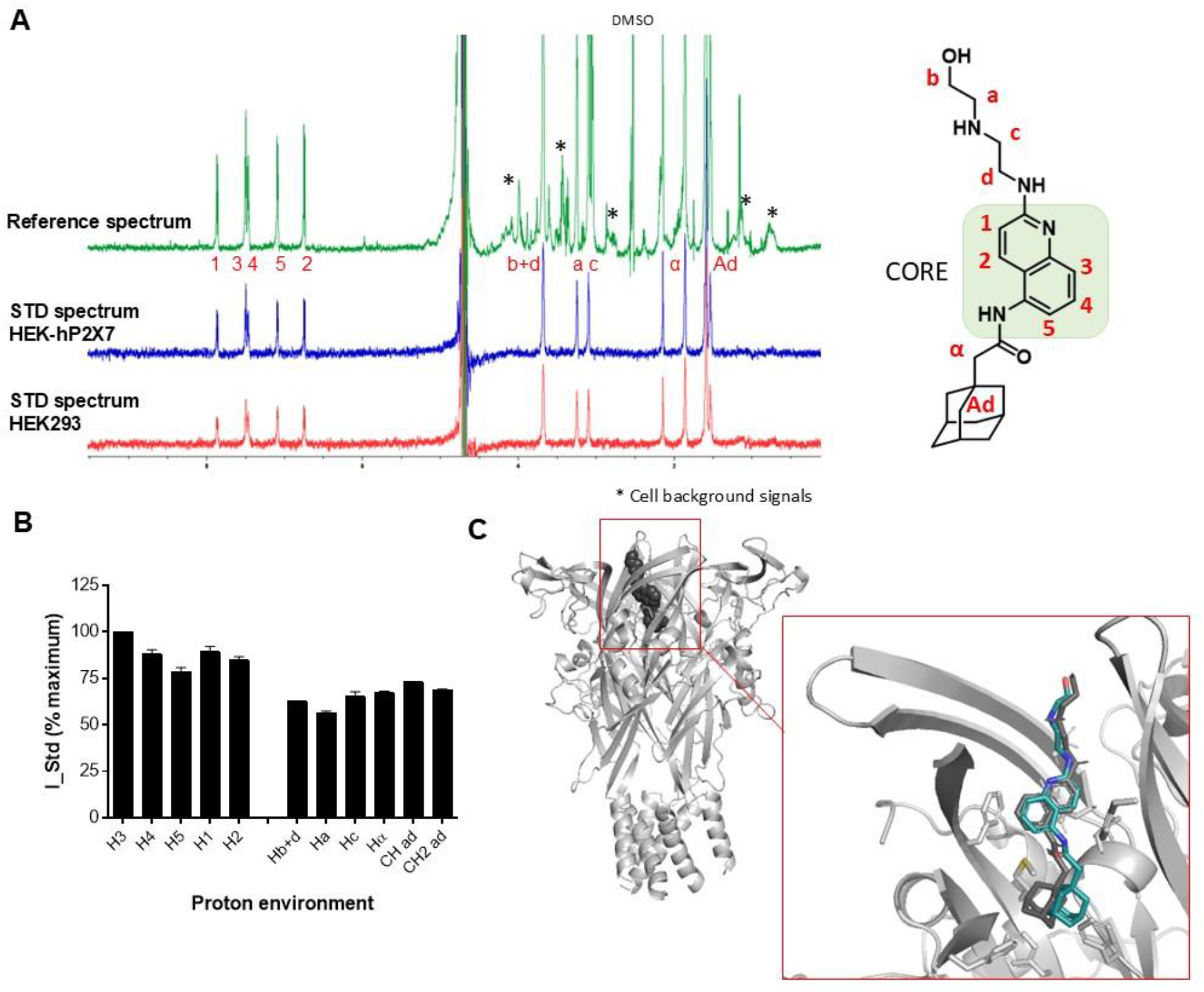
Binding profile of AZ10606120 and human P2X7 in living cells by STD NMR. **A:** unprocessed STD NMR spectra for AZ10606120; proton assignment and atom naming for AZ10606120 are shown. * indicates background signals from cells and DMSO peak is labelled. **B:** histogram reporting the normalised binding epitope for the interaction of AZ10606120 with human P2X7. Error bars represent standard deviation of 2 replicates. **C:** comparison of the docking pose for AZ10606120 to hP2X7 homology model (in grey) with the X-ray structure with pdP2X7 (in cyan) (pdb: 5u1w).

We performed the same experiments for JNJ-47965567 binding to hP2X7 and report the ligand binding epitope derived from raw NMR data (Figure 3A, 3B). The STD NMR binding epitope analysis shows that the core region for JNJ-47965567 involves ring B and the sulfur linker to ring A. We also compared the crystal structure of JNJ-47965567 at pdP2X7 (pdb:5u1x) and the computational docking of JNJ-47965567 to a human P2X7 homology model (Figure 3C).

**Figure 3:**
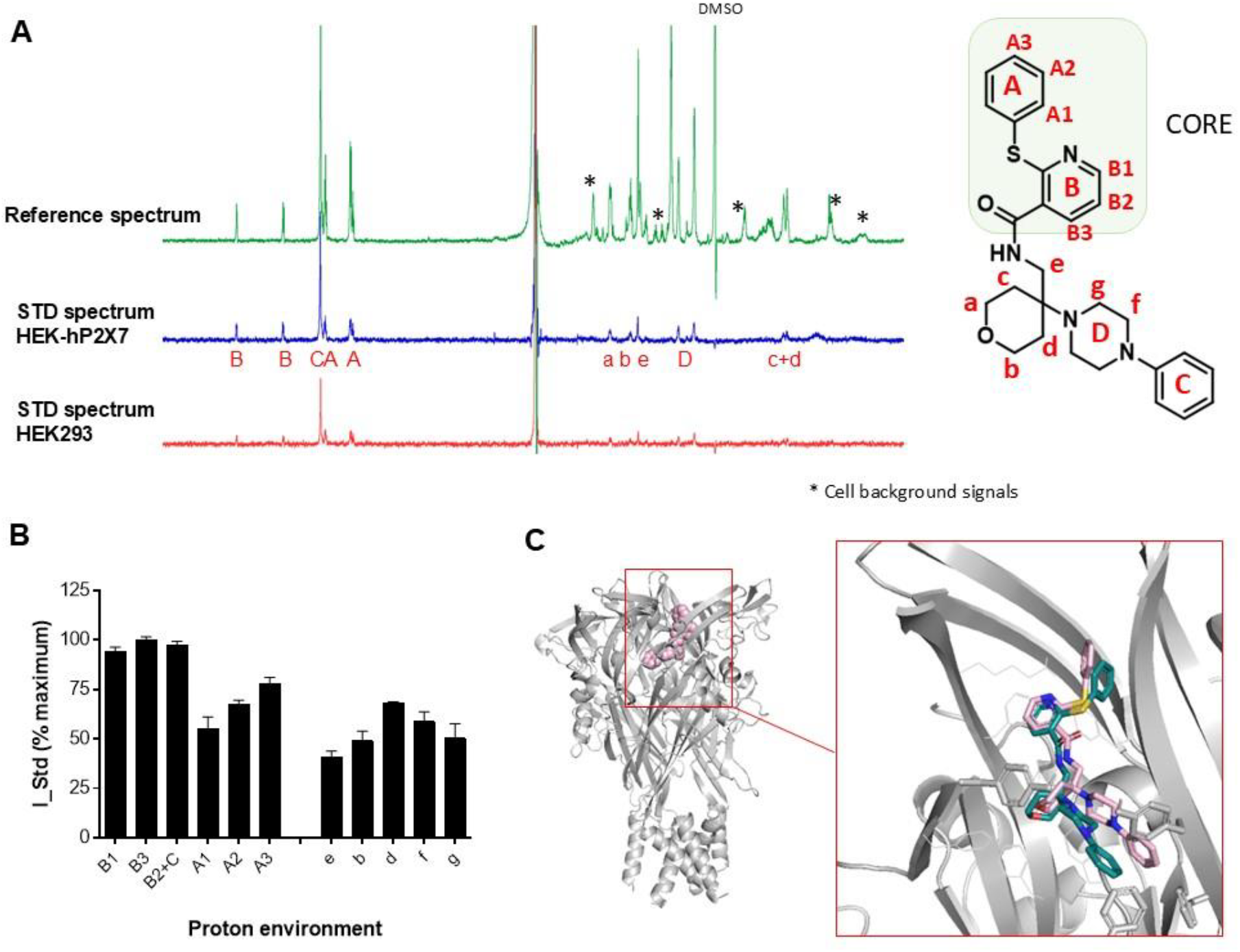
Binding profile of JNJ-47965567 and human P2X7 in living cells by STD NMR. **A:** unprocessed STD NMR spectra for JNJ-47965567, proton assignment and atom nomenclature. * indicates background signals from cells and DMSO peak is labelled. **B:** histogram reporting the normalised binding epitope for JNJ-47965567. Error bars represent standard deviation of 2 replicates **C:** comparison of the best docking pose for AZ10606120 to hP2X7 (in pink) with the X-ray structure with pdP2X7 (in cyan) (pdb: 5u1x).

When superimposing the docking poses for the two antagonists AZ10606120 and JNJ-47965567 (as well as the two pdP2X7 crystal structures, pdb: 5u1x, 5u1w), their core regions overlap (Supplementary Figure 1). For JNJ-47965567, the C-ring is deepest in the pocket (Figure 2C) and picks up very high saturation transfer suggesting close proximity to the protein surface. The tetrahydropyran ring overlaps with the space occupied by the adamantane ring in AZ10606120 and has a weaker saturation transfer. We used RedMat to correlate the experimental on-cell STD NMR data for JNJ-47965567 to the predicted computational pose and the calculated R-NOE value was 0.34, indicating a reasonable agreement.

### Pharmacology, on-cell STD NMR and docking studies on AZ10606120 and JNJ-47965567 binding to P2X7 across species (human, mouse, rat)

Once we established the ligand binding epitopes for AZ10606120 and JNJ-47965567 for their interactions with human P2X7, we explored species differences between human and rodent P2X7 receptors. By measuring intracellular calcium responses, we confirmed that agonist ATP has a similar effect at all three (Figure 4A). AZ10606120 is more potent at hP2X7 and has the weakest potency at mP2X7, while rP2X7 is intermediate (Figure 4B). IC_50_ values for both antagonists are reported in Table 1.

**Table 1:** LogIC_50_ values for P2X7 antagonists derived from fura-2 assay on stable cell lines. Standard error is indicated and mean IC_50_ shown in brackets. From n=3 independent experiments.

|  | AZ10606120 (nM) | JNJ-47965567 (nM) |
| --- | --- | --- |
| Human P2X7 | -8.10 ± 0.14 (7.9nM) | -8.04 ± 0.10 (9.0nM) |
| Rat P2X7 | -7.28 ± 0.14 (53.1nM) | -7.76 ± 0.08 (17.6nM) |
| Mouse P2X7 | -6.28 ± 0.15 (523.6nM) | -6.90 ± 0.12 (125.4nM) |

**Figure 4:**
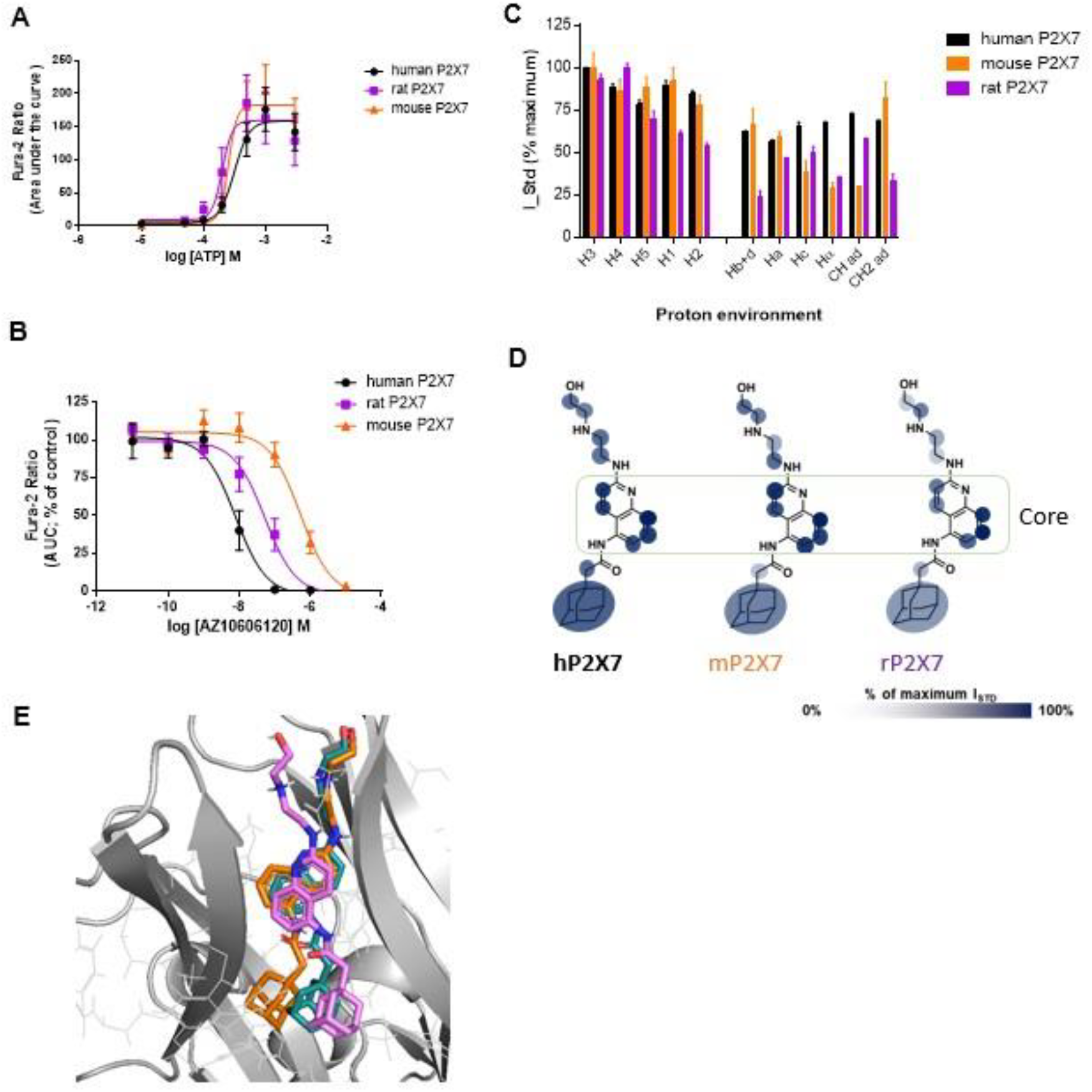
Pharmacology and binding profile of AZ10606120 interacting with rat and mouse P2X7 in living cells by STD NMR. **A:** Concentration-response curve for ATP at P2X7 orthologues; human, rat and mouse expressed in HEK-293 cells from a fura-2 calcium influx assay. **B:** Dose inhibition for AZ10606120 at P2X7 orthologues. **C:** histogram reporting the normalised binding epitope for AZ10606120 at human (black), rat (purple) and mouse (orange) P2X7. **D:** Graphical representation of the binding epitope on the chemical structure of AZ10606120 using transparency to indicate strong and weak contacts with the receptor. **E:** comparison of the best docking pose for AZ10606120 rat (magenta) and mouse (orange) with the X-ray structure with pdP2X7 (in cyan) (pdb: 5u1x).

We acquired ligand binding epitopes by STD NMR for AZ10606120 interacting with rP2X7 and mP2X7 (Figure 4C and 4D). Comparing these binding epitopes to the binding epitope at hP2X7, we observe very minor differences in the “core” aromatic region of AZ10606120. This region makes the strongest contact with the protein and this feature is conserved across the orthologues. On the contrary, the epitopes of both rP2X7 and mP2X7 AZ10606120 show lower saturation transfer to the aliphatic tail (“Hc”) and to the adamantane moiety (“Ad”) (Figure 4C, 4D). With the biggest potency difference seen between human and mouse P2X7, these results suggest that 3 ligand contact points are responsible (“Hα” “CH ad” and “Hc”) for the pharmacological difference.

Computational docking of AZ10606120 into the mP2X7 homology model and rP2X7 structure (pdb:6u9v) show similar binding poses to the pdP2X7 crystal structure (Figure 4E). Here again, in agreement with the binding epitopes, the orientation and position of the core aromatic region is conserved, while the adamantane and tail moieties have slight shifts from the human and panda P2X7. Using RedMat to correlate the experimental on-cell STD NMR data to the predicted computational poses, we achieved R-NOE values for AZ10606120 of 0.34 (mouse) and 0.54 (rat). This latter value indicates that the computational model does not represent a good correlation with the experimental NMR data suggesting there are differences in how AZ10606120 engages with the NAM pocket in rP2X7.

We then performed the same analysis for the second antagonist: from intracellular calcium measurements we determined that JNJ-47965567 had the highest potency at human P2X7 with the same rank order of potency as AZ10606120 (human>rat>mouse) (Figure 5A and Table 1), although the difference in IC_50_ values was much smaller here than for AZ10606120. Again, we performed computational docking of JNJ-47965567 into the mP2X7 homology model and rP2X7 structure (pdb:6u9v). The predicted binding orientation and position of JNJ-47965567 is similar, with differences seen in rotation of ring A and placement of ring C (Figure 5D). From the on-cell STD NMR data we can detect subtle changes in the ligand interactivity profile (Figure 5B) which must underlie the differences in pharmacological effect. The proton assigned with maximal saturation (100%) is changed from “B3” (human) to “B2+C” (rat/mouse). For both “B1” and “B3” we observed lower saturation transfer suggestive of an altered engagement between the core and the protein surface. Increases in saturation transfer to “e” “f” and “g” agree with the predicted change in placement of rings C and D (Figure 5). Using RedMat to correlate the experimental on-cell STD NMR data to the predicted computational poses, we achieved R-NOE values for JNJ-47965567 of 0.36 (mouse) and 0.29 (rat). Using the recent cryoEM model of JNJ-47965567 [23] complexed with rat P2X7 (pdb 8TRB) the R-NOE value was 0.31 suggesting a good agreement between the cryoEM structure and the experimental data.

**Figure 5:**
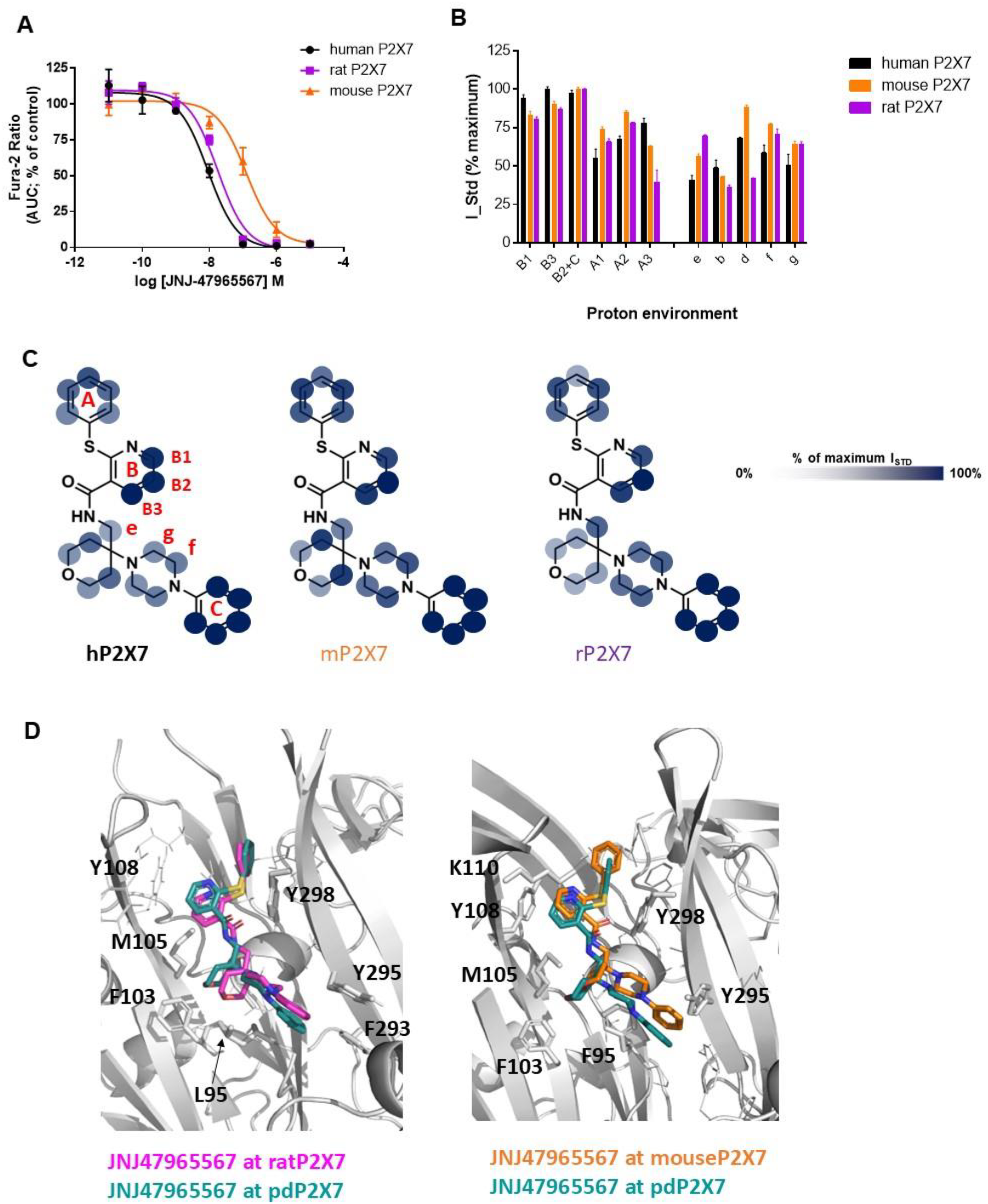
Pharmacology and binding profile of JNJ-47965567 interacting with rat and mouse P2X7 in living cells by STD NMR. **A:** Dose inhibition for JNJ-47965567 at P2X7 orthologues. **B:** histogram reporting the normalised binding epitope for JNJ-47965567 at human (black), rat (purple) and mouse (orange) P2X7. **C:** Graphical representation of the binding epitope on the chemical structure of AZ10606120 using transparency to indicate strong and weak binding. **D:** comparison of the best docking pose for AZ10606120 rat (magenta) and mouse (orange) with the X-ray structure with pdP2X7 (in cyan) (pdb: 5u1w). Key residues with side chains displayed are numbered.

### Pharmacology, on-cell STD NMR and docking studies on AZ10606120 and JNJ-47965567 binding to P2X7 across NAM-site mutants

After determining the ligand binding epitopes for AZ10606120 and JNJ-47965567 across orthologues, we explored differences across hP2X7 and selected NAM-site mutants. We chose to study three signature mutants M105A, F88A and F103A, as they have been reported to affect potency of multiple NAMs [5, 12]. We performed pharmacology studies with our overexpressed NAM-site mutant hP2X7 cell lines, including concentration response experiments to agonists ATP and BzATP to show that mutants were functional (Figure 6A, 6B). Dose inhibition curves show the effect on AZ10606120 potency as the NAM site is altered (Figure 6C) and IC_50_ values are reported in Table 2. We confirmed a loss in potency for both antagonists from hP2X7 to the three mutants. Namely, F88A and M105A show a similar loss in potency with IC_50_ dropping to ∼300 nM (from 6.8 nM), while F103A shows a major 1000-fold loss in potency. Figure 6D-E shows subtle changes in the AZ10606120 binding epitopes between hP2X7 and the NAM-site mutants. The proton assigned with maximal saturation (100%) is changed from “H3” (human wild-type) to “H5” (F88A) or “H1” (F103A) suggestive of an altered engagement between the core and the protein surface of the mutant P2X7. Indeed, mutant F103A shows the most variation in saturation transfer to the core region compared to WT human P2X7 and has additional differences in both the aliphatic tail and adamantane moieties (Figure 6D) underpinning the large difference in pharmacological potency for this antagonist.

**Table 2:** LogIC_50_ values for P2X7 antagonists at NAM-site mutants derived from fura-2 assay with standard error. Mean IC_50_ stated in brackets. From n=3 independent experiments.

|  | AZ10606120 (nM) |  | JNJ-47965567 (nM) |  |
| --- | --- | --- | --- | --- |
|  | **ATP** | **BzATP** | **ATP** | **BzATP** |
| WT hP2X7 | -8.16 ± 0.14<br>(6.8nM) | -7.65 ± 0.13<br>(22.5nM) | -7.926 ± 0.08<br>(11.7nM) | -7.59 ± 0.13<br>(26.0nM) |
| F88A hP2X7 | -6.41 ± 0.16<br>(392nM) | -6.55 ± 0.16<br>(281nM) | -6.917 ± 0.10<br>(125.3nM) | -7.27 ± 0.20<br>(53.8nM) |
| M105A<br>hP2X7 | -6.58 ± 0.34<br>(266nM) | -6.20 ± 0.21<br>(636nM) | -7.341 ± 0.28<br>(45.0nM) | -7.54 ± 0.29<br>(28.7nM) |
| F103A hP2X7 | -5.21 ± 0.54<br>(6170nM) | -5.44 ± 0.18<br>(3599nM) | -6.81 ± 0.17<br>(161.1nM) | -7.23 ± 0.10<br>(59.1nM) |

**Figure 6:**
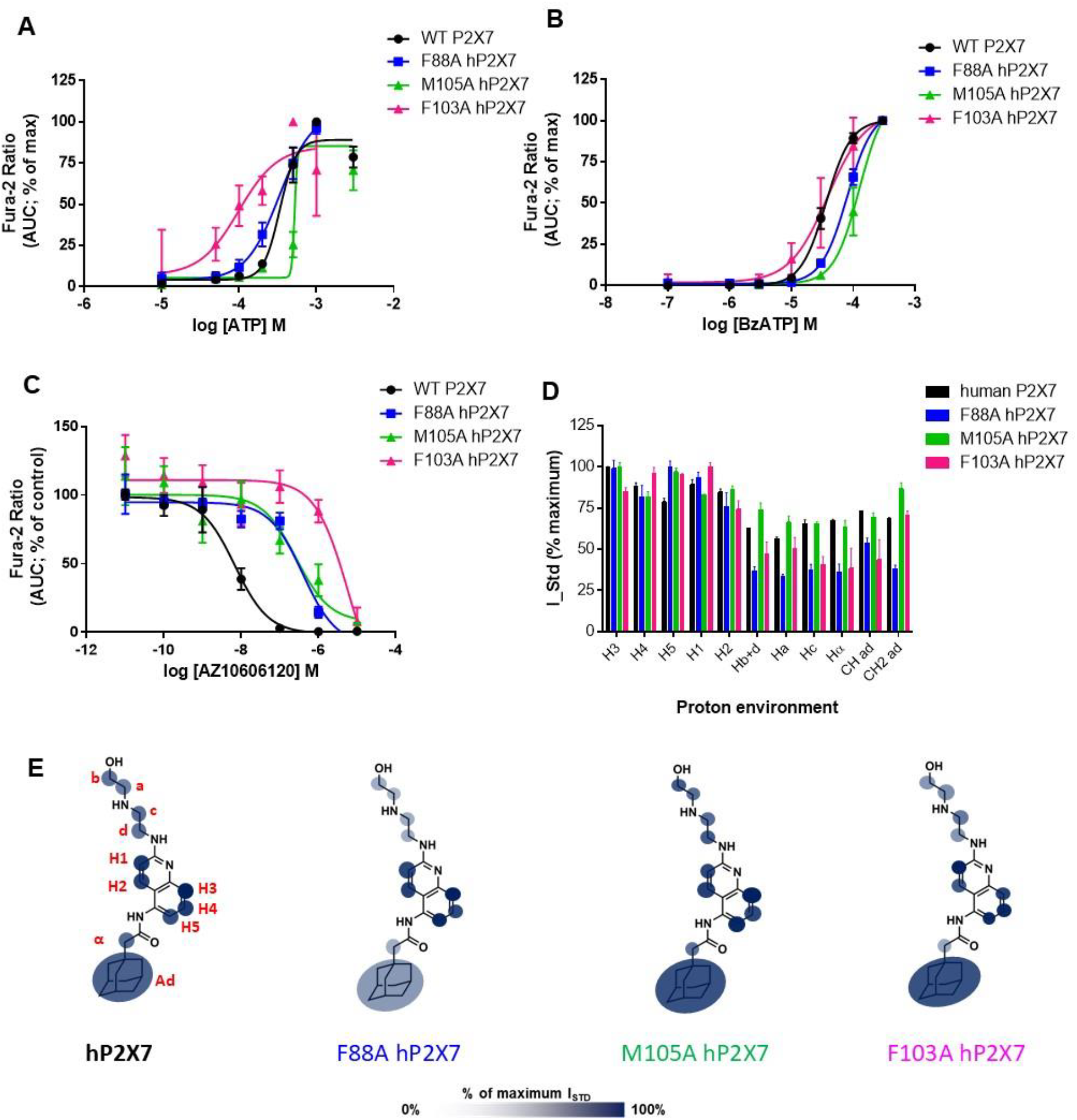
Pharmacology and binding profile of AZ10606120 interacting with WT hP2X7 and with P2X7 mutants M105A, F88A, and F103A. **A:** concentration-response curves to ATP and **B:** concentration-response curves for BzATP for the mutant hP2X7. **C:** dose inhibition curve for AZ10606120. **D:** ligand binding epitopes for AZ10606120 in histogram form. Error bars represent standard deviation from 2 replicates. **E:** Graphical representation of the binding epitope on the chemical structure of AZ10606120 using transparency to indicate strong and weak binding.

We performed computational docking for the mutants (see Supplementary Figure 2) and no dramatic variation from the binding mode at the wild-type hP2X7 was observed. RedMat analysis of these poses against the STD NMR data, reports R-NOE value,s of 0.29 (M105A), 0.38 (F88A) and 0.26 (F103A), validating the models of M105A- and F103A-ligand complexes. The STD NMR data therefore gives unique insights into how the interactivity profile between the ligand and the protein changes under experimental conditions.

Figure 7 shows the pharmacology and structural analysis for JNJ-47665567, with IC_50_ values reported in Table 2. Similar to the species differences, we observed a much smaller loss in potency between hP2X7 and the NAM-site mutants for JNJ-47965567, with only a 10-fold increase in IC_50_ for F88A and F103A, and a less than 5-fold increase for M105A-hP2X7. From the on-cell STD NMR data we can detect changes in the ligand interactivity profile (Figure 5B) which must underlie the differences in pharmacological effect. The proton assigned with maximal saturation (100%) is changed from “B3” (human) to “B1” (mutants) and there is a considerable decrease in saturation transfer to “B3” and “B2+C” suggestive of an altered engagement between the core region and the protein surface. Reductions in saturation transfer to “d” “f” and “g” are also seen with F88A and F103A which correlate with the biggest pharmacological difference seen.

**Figure 7:**
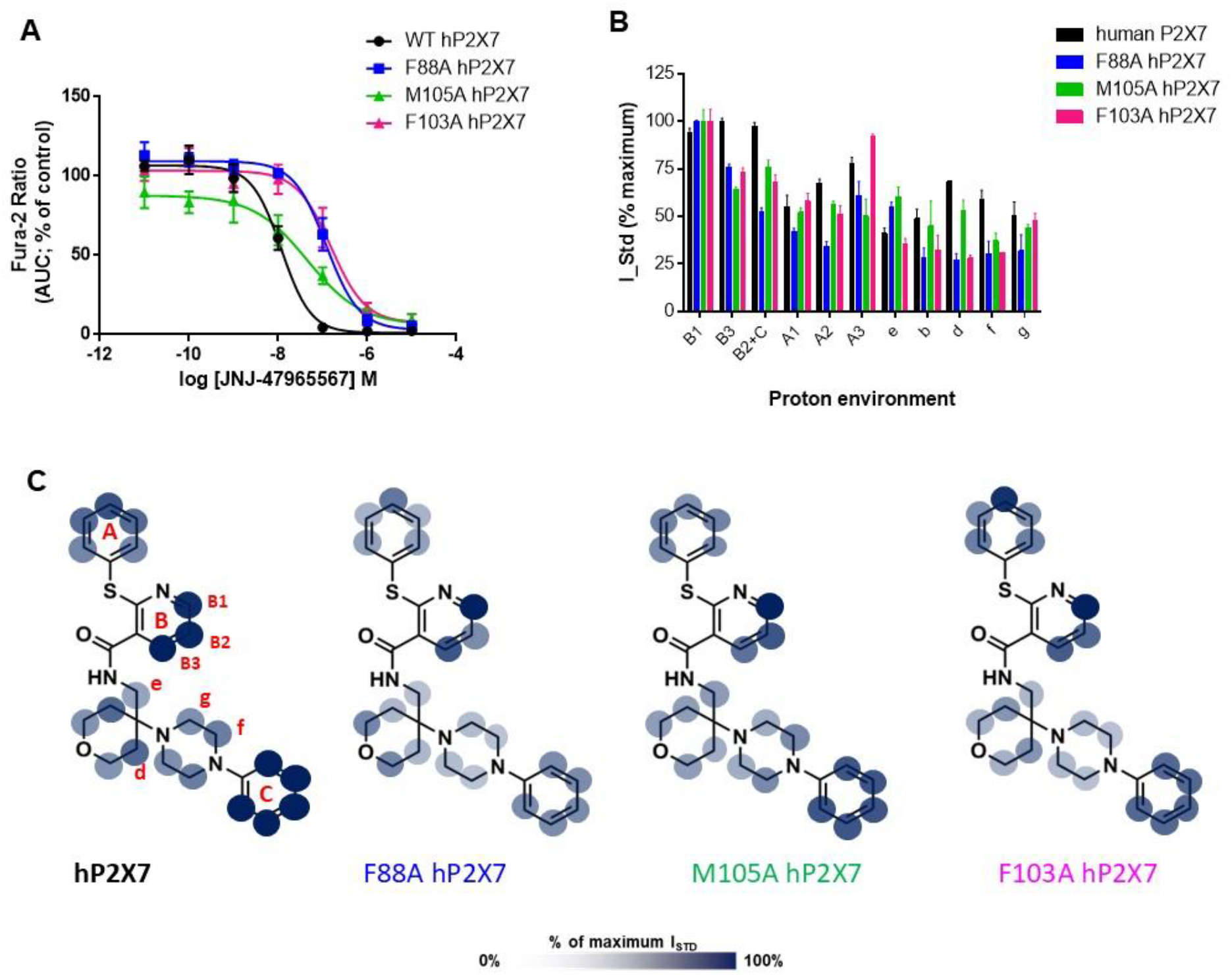
Pharmacology and binding profile of JNJ-47965567 interacting with WT hP2X7 and with P2X7 mutants M105A, F88A, and F103A. **A:** dose inhibition curve for JNJ-47965567. **B:** ligand binding epitopes for JNJ-47965567 in histogram form. Error bars represent standard deviation of 2 replicates. **C:** Graphical representation of the binding epitope on the chemical structure of JNJ-47965567 using transparency to indicate strong and weak binding.

The computational docking did not predict dramatic variations in the binding pose of JNJ-47965567 (Supplementary Figure 3), and from RedMat analysis, these poses do not correlate well with the on-cell STD NMR data. RedMat reports R-NOE values of 0.35 (M105A-hP2X7), 0.51 (F103A-hP2X7) and 0.57 (F88A-hP2X7) indicating that the computational prediction is likely to be inaccurate for F88A and F103A. However, it is clear that the STD NMR data gives unique insights into how the interactivity profile between the ligand and the protein changes under experimental conditions.

## Discussion

### Comparison of binding mode and potency across species

With the publication of the crystal structure of pdP2X7 in complex with five different negative allosteric modulators [5], a structural model of how such ligands engage with a mammalian P2X7 was first visualised. Overlaying these ligands can yield some information regarding key chemical groups and looking at potential interactions can yield information about key interacting residues. Here, we provide for the first time, experimental evidence that the overlapping “core” regions of AZ10606120 and JNJ-47965567 make the strongest contacts with the P2X7 protein membrane-embedded on living cells. The cores are close to an aromatic-rich region of the NAM binding pocket composed of phenylalanine residues (88, 108) and tyrosine 298. This suggests that aromatic-aromatic interactions could be a main factor for the high affinity interactions of these two ligands to the hP2X7 NAM binding pocket. JNJ-47965567 is more tightly bound into the NAM pocket with additional strong contacts being made with the deepest part of the molecule, the phenyl moiety on the piperazine ring (C-ring in Figure 3). Our on-cell STD NMR binding epitope mapping was corroborated by the docking poses obtained for the two ligands using a hP2X7 homology model and a template-based docking approach. The docking pose predicted is highly similar to the crystal structure of pdP2X7 in complex with AZ10606120 and JNJ-67965567 and is experimentally validated through calculation of a reduced relaxation matrix (RedMat) to predict the theoretical binding epitope maps from 3D molecular models of the protein-ligand complexes [22].

Our investigations on P2X7 rodent homologues suggest that the central core of AZ10606120 makes interactions with the NAM pocket across species, whereas saturation transfer to the aliphatic tail and adamantane moiety of AZ10606120 are weaker for the rodent P2X7 receptors. A key difference between mouse P2X7 and human P2X7 NAM-site is at amino acid 312, which is a valine in human, but an alanine in mouse/rat. This substitution would cause the rodent NAM pocket to be slightly bigger, accounting for some loss of STD intensities for the adamantane moiety of AZ10606120 (Figure 4D). An additional difference between rat P2X7 and human P2X7 is amino acid 95 – which is phenylalanine in human/mouse and leucine in rat [24]. This residue sits at the very base of the NAM pocket and is also expected to make interactions with the adamantane moiety of AZ10606120. In the absence of F95, the rat NAM pocket would be considerably bigger than the human and mouse P2X7 NAM-site explaining the further loss of STD signal on the adamantane moiety. It is likely that, in the absence of the bulky F95, the AZ10606120 ligand has a different orientation, but it cannot extend deeper into the pocket due to the core region interactions. Indeed, the docking poses confirm the hypothesis that the core region is “anchoring” the ligand at a fixed depth through aromatic-aromatic interactions in the lipophilic binding pocket, giving very little freedom for the ligand to rearrange and occupy the larger pockets in a more effective way. This can be mainly ascribed to the most rigid and linear character of AZ10606120, so that once it is “anchored” it cannot vary its orientation to occupy the space differently. This also implies that this ligand retains a similar position from the smaller human P2X7 NAM pocket to the larger rodent P2X7 NAM pocket. The higher R-NOE value (0.54) for the docking generated model of AZ10606120 at rP2X7 suggests that the docking pose for this complex is not representative of the binding mode of AZ10606120 in the much larger rat P2X7 NAM pocket. This is difficult to reconcile using computational docking, and more structural data (x-ray crystallography or cryoEM) would be required to clarify this.

Looking closely at the biggest pharmacological difference for AZ10606120 (which is between human and mouse P2X7) to correlate this with the on-cell STD NMR data, we suggest that 3 major contact points could be responsible for this change in potency - “Hα” “CH ad” and “Hc”. This information could be used in a drug discovery program to improve the molecular fit to the rodent P2X7 NAM site, where further chemical modifications could involve the aliphatic tail and adamantane moieties. This highlights the utility of this approach in analysing a library of ligands during structure-activity relationship investigations.

For JNJ-47465567 the core region also receives high saturation transfer, a feature that is somewhat conserved across species, suggesting tight anchoring to the aromatic-rich region in the centre of the P2X7 NAM binding pocket. We did observe that “B1” and “B3” showed lower saturation transfer suggestive of an altered engagement between the core and the protein surface with increases in saturation transfer to “e” “f” and “g” protons also observed (Figure 5). The variability in pharmacological potency of JNJ-47965567 across species was less than for AZ10606120 with only a 13-fold difference between hP2X7 and mP2X7 IC_50_ values.

However, we observed more variation in overall saturation transfer to JNJ-47965567 between hP2X7, mP2X7 and rP2X7, mostly seen at the tetrahydropyran – piperazine region. The tetrahydropyran ring of JNJ-47965567 overlaps with the adamantane moiety of AZ10606120 (Supplementary Figure 1), and in hP2X7 the adamantane moiety averages 70% STD intensity whereas the tetrahydropyran ring has 50-70% STD intensities. Therefore, we see clear consistencies between the two ligands in the hP2X7 NAM pocket. With both the “core” and the deepest phenyl moiety of JNJ-47965567 being well anchored within the NAM pocket, this explains why there is little difference in pharmacological effect with this ligand (confirming that seen in [25]) even though there is some variation in the architecture of NAM pocket across species. Certainly, the detailed ligand binding epitope provided by the on-cell STD NMR approach is of great relevance to identify the regions available for chemical modification, for a ligand to undergo further improvement.

### Comparison of binding mode and potency across mutants

Pharmacology and binding epitope analysis were performed for three hP2X7 mutants, namely M105A, F88A and F103A, for both AZ10606120 and JNJ47465567 ligands. As expected from other studies [12], we did observe a reduction in potency relative to the WT hP2X7 for the specific mutants selected and this reduction was much more pronounced for AZ10606120 than for JNJ-47465567. As observed for the P2X7 orthologues, there was variation in binding epitopes for AZ10606120, some of which was again related to the aliphatic tail and adamantane moieties. We did observe altered saturation transfer to the core region of AZ10606120 with a switching of proton assigned with maximal saturation (100%) from “H3” (human) to “H5” (F88A) or “H1” (F103A). Focusing on F103A, these alterations in binding epitope are responsible for the large difference in pharmacological potency for this antagonist and this technique is useful for highlighting how this mutation affects the antagonist interactivity with the P2X7 NAM-site surface.

With JNJ-47965567, there were similar alterations in the saturation transfer to the core region in mutants exhibiting the largest change in pharmacological potency (F88A, F103A). RedMat did not validate the poses at F88A-hP2X7 and F103A-hP2X7 against the STD NMR binding epitope data, suggesting that the crystal structure pdP2X7-like orientation of this ligand is not valid for these NAM-site mutants. In these cases, it may be that there are major alterations/rearrangements of the aromatic networks within the NAM pocket which allows JNJ-47965567 to interact differently. This hypothesis is compatible with the reduction in the STD intensities associated with the deepest phenyl moiety (C-ring) for F88A-hP2X7 and F103A-hP2X7 plus the changes in the core region.

In conclusion, here we show for the first time that saturation transfer difference (STD) NMR can be used on a mammalian cell line over-expressing a membrane-embedded ion channel to gain structural insights on receptor-ligand interactions, in their native environment. We performed on-cell STD NMR for the structural investigation of NAM antagonists as bound to P2X7 ion channels in their physiological membrane-bound environment. The combination of STD NMR with molecular docking has been shown to be an insightful approach providing the first NMR-validated 3D models for hP2X7 and mP2X7 as bound to AZ10606120 and JNJ-47465567, and rP2X7 as bound to JNJ-47465567. The inexpensive and versatile nature of NMR and STD NMR relative to x-ray crystallography and cryo-EM for membrane-embedded ion channels should be highlighted, in addition to the capability of studying these biological systems in solution and in a native environment. We have shown the potential of on-cell STD NMR to pick up on differences in the binding mode upon binding pocket variation and, with some limitations, we have been able to correlate this to pharmacology data, paving the way for a wider applicability to structure-activity-relationship (SAR) studies for receptors embedded in live cells. This approach will be extremely useful for the design of new and successful drugs targeting ion channels and other embedded receptors and we envisage this approach to become a new frontier in drug discovery.

## Acknowledgements

We thank the University of East Anglia Faculty of Science NMR platform for the use of the 800MHz spectrometer. This work was supported by UKRI Future Leaders Fellowship to MW (grant number MR/T044020/1) and UEA School of Pharmacy.

## References

1. Deussing, J.M. and E. Arzt, P2X7 Receptor: A Potential Therapeutic Target for Depression? Trends in Molecular Medicine, 2018. 24(9): p. 736–747.

2. Horváth, G., et al., P2X7 Receptors Drive Poly(I:C) Induced Autism-like Behavior in Mice. The Journal of Neuroscience, 2019. 39(13): p. 2542.

3. Szabó, D., et al., Maternal P2X7 receptor inhibition prevents autism-like phenotype in male mouse offspring through the NLRP3-IL-1β pathway. Brain, Behavior, and Immunity, 2022. 101: p. 318–332.

4. Lara, R., et al., P2X7 in Cancer: From Molecular Mechanisms to Therapeutics. Frontiers in Pharmacology, 2020. 11.

5. Karasawa, A. and T. Kawate, Structural basis for subtype-specific inhibition of the P2X7 receptor. eLife, 2016. 5: p. e22153.

6. Dane, C., L. Stokes, and W.T. Jorgensen, P2X receptor antagonists and their potential as therapeutics: a patent review (2010–2021). Expert Opinion on Therapeutic Patents, 2022. 32(7): p. 769–790.

7. Eser, A., et al., Safety and Efficacy of an Oral Inhibitor of the Purinergic Receptor P2X7 in Adult Patients with Moderately to Severely Active Crohn’s Disease: A Randomized Placebo-controlled, Double-blind, Phase IIa Study. Inflammatory Bowel Diseases, 2015. 21(10): p. 2247–2253.

8. Keystone, E.C., et al., Clinical evaluation of the efficacy of the P2X7 purinergic receptor antagonist AZD9056 on the signs and symptoms of rheumatoid arthritis in patients with active disease despite treatment with methotrexate or sulphasalazine. Annals of the Rheumatic Diseases, 2012. 71(10): p. 1630.

9. Stock, T.C., et al., Efficacy and Safety of CE-224,535, an Antagonist of P2X7 Receptor, in Treatment of Patients with Rheumatoid Arthritis Inadequately Controlled by Methotrexate. The Journal of Rheumatology, 2012. 39(4): p. 720.

10. Ali, Z., et al., Pharmacokinetic and pharmacodynamic profiling of a P2X7 receptor allosteric modulator GSK1482160 in healthy human subjects. British Journal of Clinical Pharmacology, 2013. 75(1): p. 197–207.

11. Recourt, K., et al., Characterization of the central nervous system penetrant and selective purine P2X7 receptor antagonist JNJ-54175446 in patients with major depressive disorder. Translational Psychiatry, 2023. 13(1): p. 266.

12. Allsopp, R.C., et al., Unique residues in the ATP gated human P2X7 receptor define a novel allosteric binding pocket for the selective antagonist AZ10606120. Scientific Reports, 2017. 7(1): p. 725.

13. Bin Dayel, A., R.J. Evans, and R. Schmid, Mapping the Site of Action of Human P2X7 Receptor Antagonists AZ11645373, Brilliant Blue G, KN-62, Calmidazolium, and ZINC58368839 to the Intersubunit Allosteric Pocket. Molecular Pharmacology, 2019. 96(3): p. 355.

14. Mayer, M. and B. Meyer, Characterization of Ligand Binding by Saturation Transfer Difference NMR Spectroscopy. Angewandte Chemie International Edition, 1999. 38(12): p. 1784–1788.

15. Mayer, M. and B. Meyer, Group Epitope Mapping by Saturation Transfer Difference NMR To Identify Segments of a Ligand in Direct Contact with a Protein Receptor. Journal of the American Chemical Society, 2001. 123(25): p. 6108–6117.

16. Angulo, J. and P.M. Nieto, STD-NMR: application to transient interactions between biomolecules—a quantitative approach. European Biophysics Journal, 2011. 40(12): p. 1357–1369.

17. Claasen, B., et al., Direct Observation of Ligand Binding to Membrane Proteins in Living Cells by a Saturation Transfer Double Difference (STDD) NMR Spectroscopy Method Shows a Significantly Higher Affinity of Integrin αIIbβ3 in Native Platelets than in Liposomes. Journal of the American Chemical Society, 2005. 127(3): p. 916–919.

18. Mari, S., et al., 1D Saturation Transfer Difference NMR Experiments on Living Cells: The DC-SIGN/Oligomannose Interaction. Angewandte Chemie International Edition, 2005. 44(2): p. 296–298.

19. Palmioli, A., et al., On-cell saturation transfer difference NMR for the identification of FimH ligands and inhibitors. Bioorganic Chemistry, 2021. 112: p. 104876.

20. Palmioli, A., et al., Multivalent calix[4]arene-based mannosylated dendrons as new FimH ligands and inhibitors. Bioorganic Chemistry, 2023. 138: p. 106613.

21. Bertuzzi, S., et al., Exploring Glycan-Lectin Interactions in Natural-Like Environments: A View Using NMR Experiments Inside Cell and on Cell Surface. Chemistry – A European Journal, 2025. 31(10): p. e202403102.

22. Nepravishta, R., et al., Fast Quantitative Validation of 3D Models of Low-Affinity Protein–Ligand Complexes by STD NMR Spectroscopy. Journal of Medicinal Chemistry, 2024. 67(12): p. 10025–10034.

23. Oken, A.C., et al., P2X7 receptors exhibit at least three modes of allosteric antagonism. Science Advances, 2024. 10(40): p. eado5084.

24. Michel, A.D., et al., Identification of regions of the P2X7 receptor that contribute to human and rat species differences in antagonist effects. British Journal of Pharmacology, 2008. 155(5): p. 738–751.

25. Bhattacharya, A., et al., Pharmacological characterization of a novel centrally permeable P2X7 receptor antagonist: JNJ-47965567. British Journal of Pharmacology, 2013. 170(3): p. 624–640.

