## Supplementary figures for "On-cell Saturation Transfer Difference (STD) NMR on ion channels: characterizing negative allosteric modulator binding interactions of P2X7"

**Monaco et al, Supplementary information**

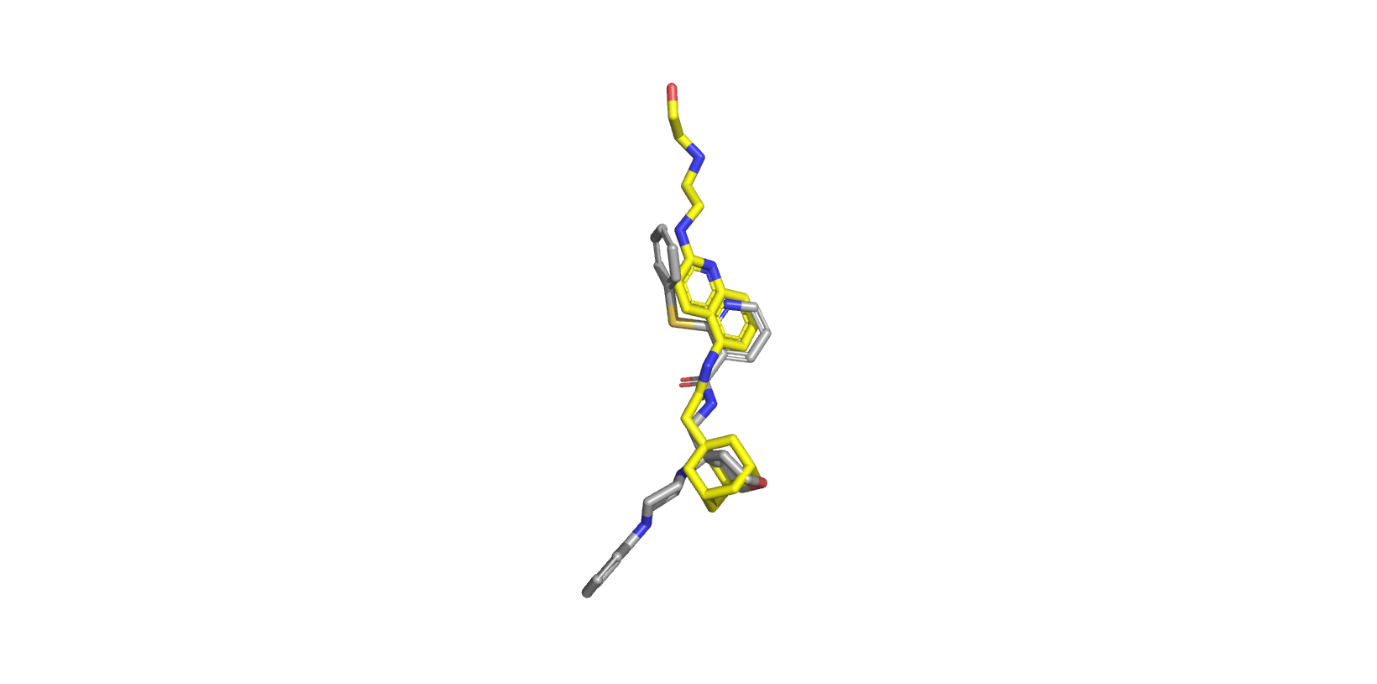

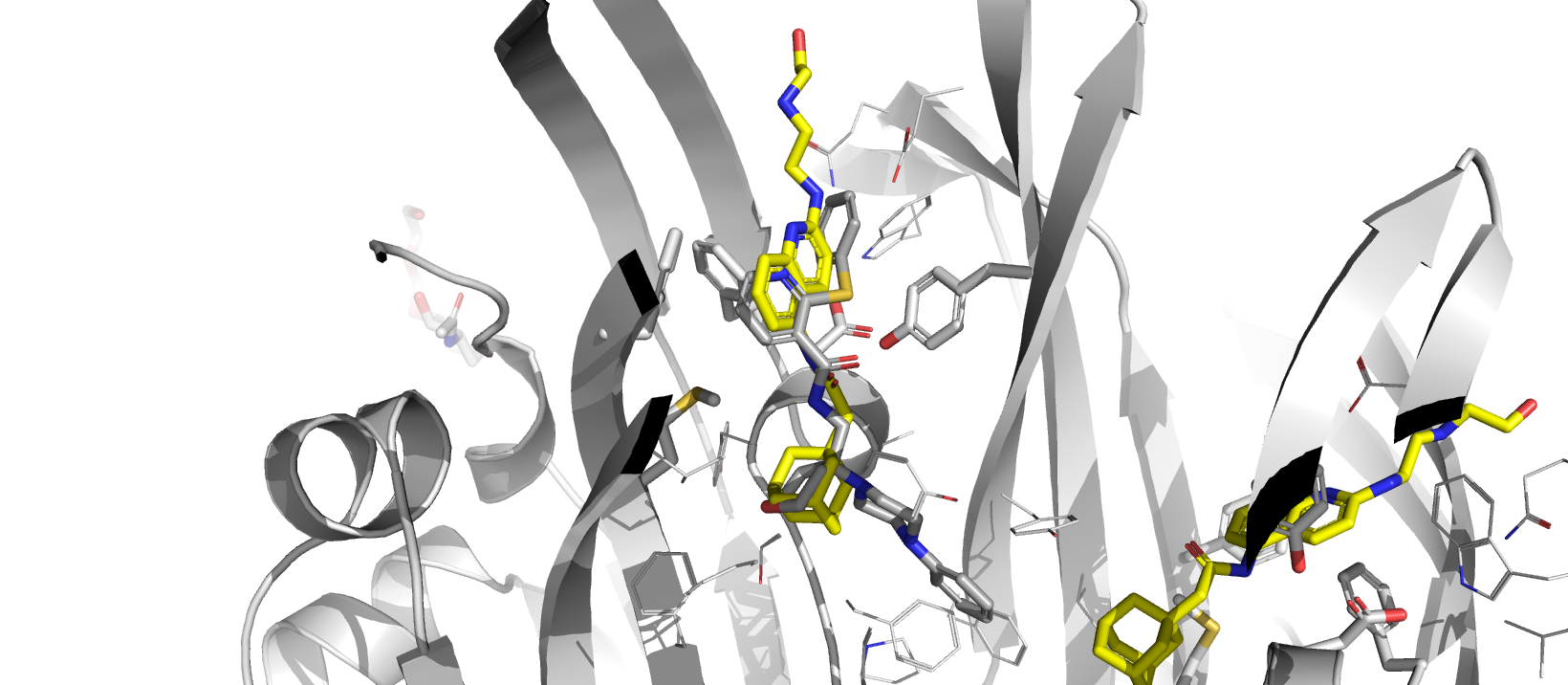

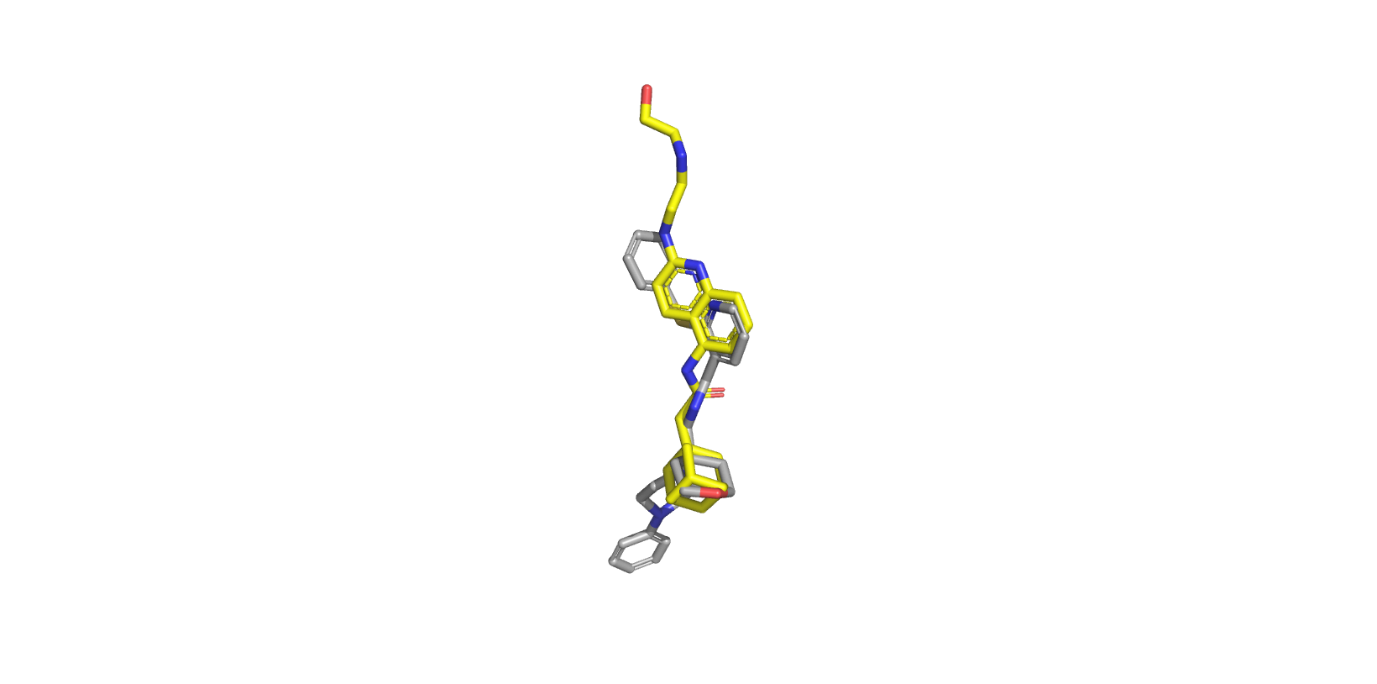

**D92**

**M105**

**F88**

**F108**

**Y298**

**Supplementary Figure 1**: Comparison of AZ10606120 (yellow) position in P2X7 NAM pocket with JNJ-47965567 (grey) position. Taken from crystal structure of pdP2X7 in complex with both ligands (pdb entries: 5u1w and 5u1x).

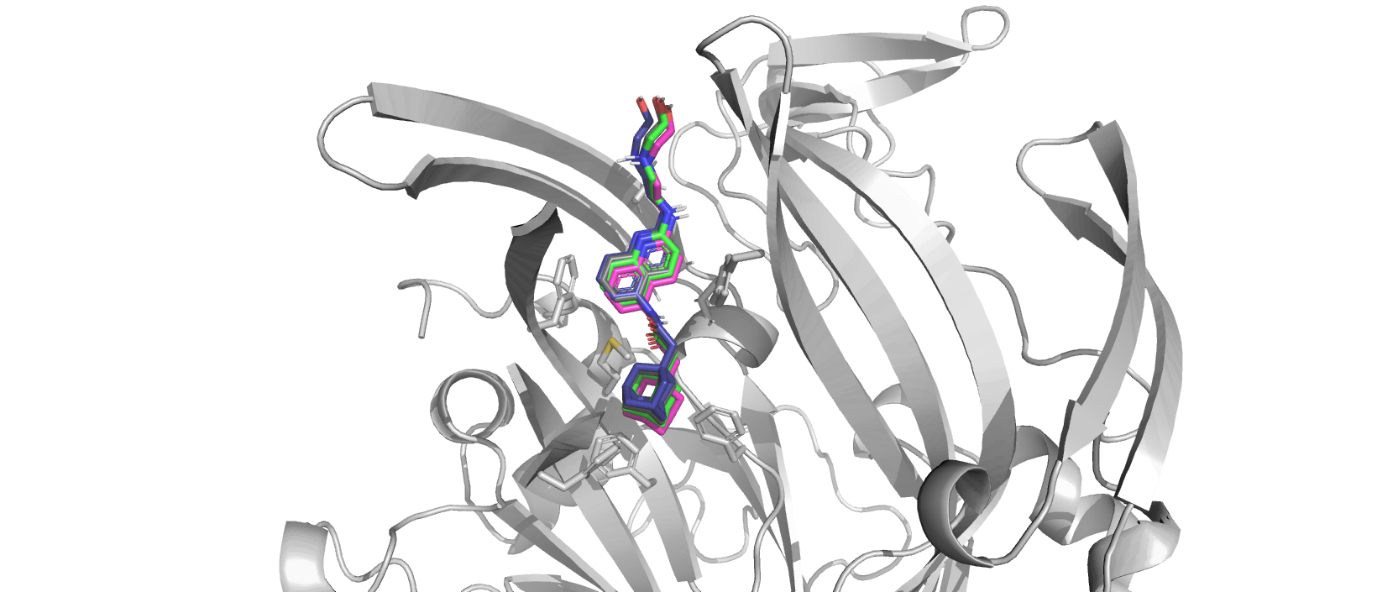

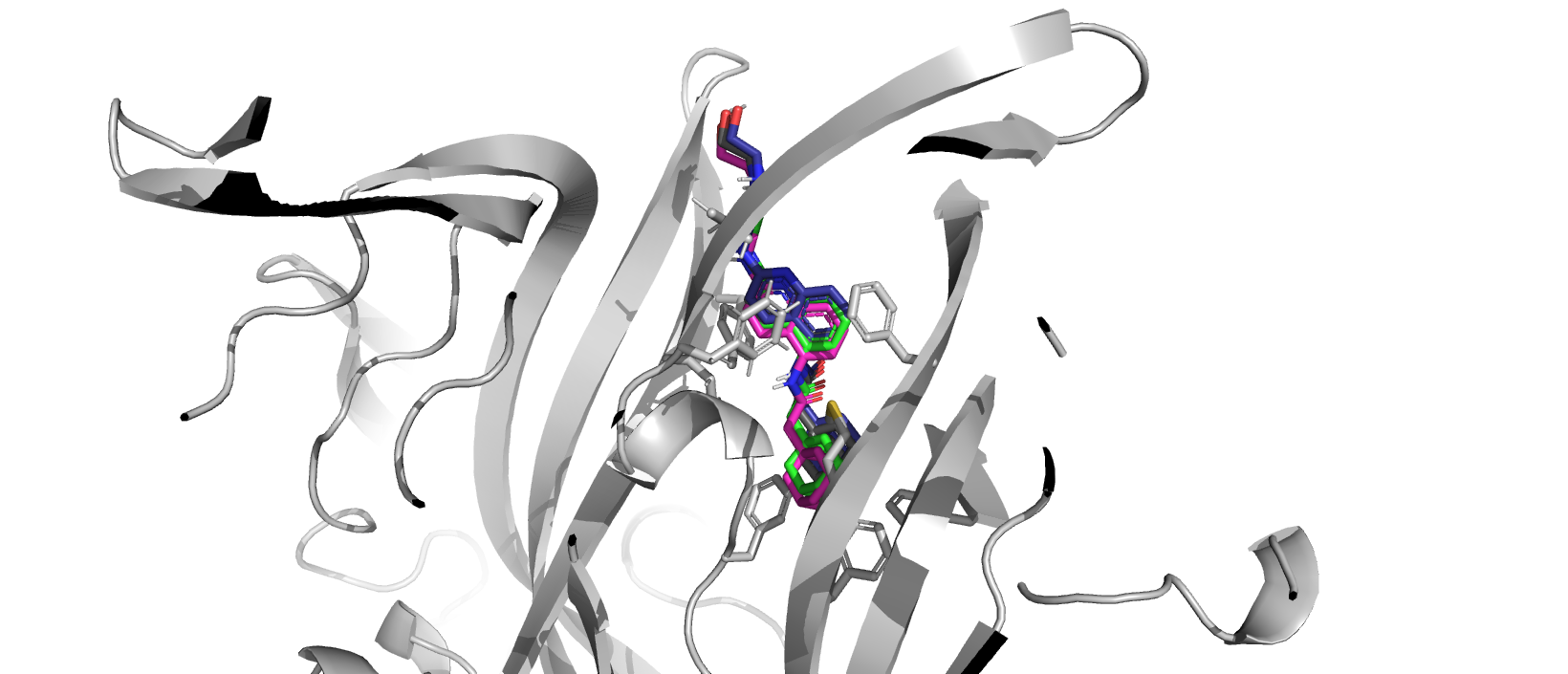

**F88**

**M105**

**F103**

**Supplementary Figure 2**: Comparison of AZ10606120 docked to human P2X7 or NAM-site mutant hP2X7 with little variation in predicted pose. F88A-hP2X7 (blue), M105A-hP2X7 (green) and F103A-hP2X7 (magenta) poses shown overlaid with hP2X7 (grey).

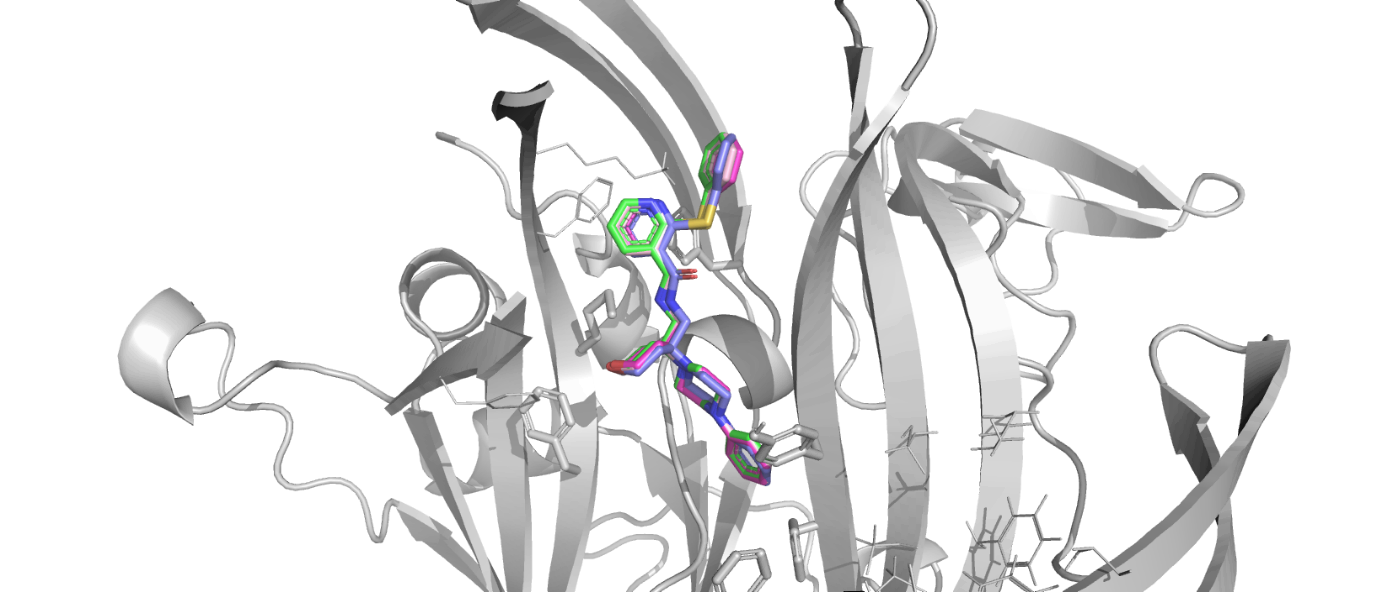

**F88**

**M105**

**F103**

**Supplementary Figure 3**: Comparison of JNJ-47965567 docked to human P2X7 or NAM-site mutant hP2X7 with little variation in predicted pose. F88A-hP2X7 (blue), M105A-hP2X7 (green) and F103A-hP2X7 (magenta) poses shown overlaid with hP2X7 (grey).
